# Multiple bursting patterns in Lateral Habenula neurons: experiments and computational model

**DOI:** 10.1101/2025.01.23.634464

**Authors:** Dmitry Fedorov, Fabien Campillo, Mathieu Desroches, Edgar Soria-Gómez, Serafim Rodrigues, Joaquin Piriz

## Abstract

The Lateral Habenula (LHb) is a small brain structure specialized in encoding aversive signals. Bursting activity in the LHb has been consistently linked to mood regulation, with increased bursting activity proposed to promote depressive behaviors. Bursting is a complex dynamic process that has been extensively studied and modeled in other neuronal contexts. However, at the LHb this type of activity has typically been described only as transient periods of high frequency firing. Here, to provide a deeper understanding of LHb bursting, we analyzed this activity from the perspective of dynamical systems. Ex vivo, LHb neurons display a variety of bursting patterns, characterized at one extreme by a dominating square-wave type and in other by parabolic type, plus transitional forms referred to as triangular bursting. Notably, these bursting patterns, which reflect different LHb output modes, can occur within the same neuron, suggesting that they may correspond to distinct dynamic states of the same LHb neuron. To capture these complex behaviors, we propose an idealized multiple-timescale dynamical model. This model successfully reproduces the three main bursting patterns observed in experimental data. Furthermore, we identify a special point in the parameter space, termed the saddle-node homoclinic bifurcation, which acts as an organizing center demarcating the boundary between the two primary bursting patterns and around which the third pattern appear. Our model suggests that LHb bursting activity is structured around distinct dynamic states with potentially diverse and unexplored impacts on mood regulation. By providing new insights into the dynamic principles underlying LHb bursting, this framework may advance our understanding of its biological significance.

## 1 Introduction

The Lateral habenula (LHb) is a highly conserved structure in the vertebrate brain, that serves as a crucial communication hub between the limbic system and the midbrain. Hyperactivity of the LHb has been implicated in several psychiatric disorders characterized by heightened aversion, particularly depression [15]. Indeed, the LHb has been proposed as a key brain target underlying the antidepressant effects of ketamine [6]. Additionally, its direct modulation through deep-brain stimulation is currently being explored as a potential treatment for depression [1].

The LHb is predominantly composed of small glutamatergic neurons with high input resistance (*>* 500 MΩ) that receive both glutamatergic and GABAergic inputs. Ex vivo, these neurons exhibit complex patterns of spontaneous spiking activity, which have been qualitatively categorized into four modes: silent, non-regular tonic spiking, regular tonic spiking and bursting [28]. However, LHb neurons can transition between these patterns, suggesting that these modes are not associated with different neuronal classes but with specific dynamic states.

Among these activity modes, silent and tonic patterns are the most prominent, while only a minority of neurons display spontaneous bursting activity. However, the majority (≈90%) of LHb neurons exhibit rebound bursting after negative current injection, indicating that bursting is a general property, although largely latent, of LHb neurons [28]. Bursting activity in the LHb has attracted significant attention due to its association with depression-like states in animal models. In these models, the proportion of LHb neurons exhibiting bursting increases. Moreover, experimentally inducing bursting can reproduce depressive-like behaviors, whereas pharmacological agents that suppress bursting, most notably ketamine, have been shown to counteract these effects [3, 6, 8, 9, 16, 19, 22, 25, 30].

However, the specific reasons why LHb bursting activity is linked to depression-like states remain unclear. Bursting activity is characterized by a temporal alternation between periods of high-frequency firing and quiescence. This implies a temporal arrangement of the neuronal output, but not necessarily an increase in overall neuronal output. Thus, from a functional perspective, it is not immediately apparent why bursting activity of the LHb contributes to depressive states. Moreover, LHb bursting activity has been primarily described in broad terms as periods of heightened firing [6, 9, 20, 30], contrasting with the detailed analyses and classifications that have been developed for bursting activity in other neural contexts [11, 17]. To fill this gap, in this article we studied and classified bursting activity at the LHb from the perspective of dynamical systems. Moreover, we present an initial phenomenological model to describe the activity of LHb neurons. From a neurodynamics perspective, we observed a gradient of bursting patterns ranging from square-wave to parabolic bursts, as well as intermediate forms referred to as triangular (see section 2.3.2 for burst type criteria). To account for this diversity of dynamical behaviors, we propose a minimal phenomenological model capable of capturing these patterns. The model possesses an organizing center in parameter space, enabling transitions between different behaviors driven by the two main slow processes at play in the system. We further conjecture that these bursting patterns arise from distinct sets of conductances, representing different modes of information transmission by LHb neurons, each with potentially unique implications on depressive-like behaviors.

## 2 Materials and methods

### 2.1 Animals

Experiments were conducted on 7 female and 17 male mice (6-9 months old) C57BL/6JRj mice (Janvier Labs, France). Animals were kept at the animal facility of the University of the Basque Country, Bizkaia Campus, under controlled temperature and humidity conditions with a 12/12 hours dark-light cycle. Food and drinking water were provided *ad libitum*. All procedures adhered the European Directive 2010/63/EU, NIH guidelines and were approved by the Ethics Committees of the University of the Basque Country EHU/UPV. (Leioa, Spain; CEEA M20/2024/213).

### 2.2 Electrophysiology

#### 2.2.1 Preparation of brain slices

Mice were anesthetized with isoflurane (IsoFlo 100%, Zoetis), and quickly decapitated into chilled and gassed NMDG based artificial cerebro spinal fluid (ACSF) of the following composition (in mM) [26]: 2.5 KCl, 1.25 NaH_2_PO_4_, 30 NaHCO_3_, 25 glucose, 20 HEPES, 5 Na-ascorbate, 2 Thiourea, 3 Na-pyruvate, 92 NMDG, 10 MgSO_4_, 0.5 CaCl_2_). The brain was extracted and sliced on a vibratome (VT1000S or VT1200S, Leica) in the same solution to a thickness of 250 µm. The slices were then recovered in 36 °C NMDG-ACSF solution for 40 minutes, with gradual introduction of Na^+^ into solution following the Na^+^ spike-in procedures described by [26]. The slices were then transferred to room temperature holding-ACSF (same as NMDG-ACSF, with 92 mM NaCl instead of NMDG, 2 mM MgSO_4_, 2 mM CaCl2) storage solution for the day.

The brain was extracted and sliced on a vibratome (VT1000S or VT1200S, Leica) in the same solution to a thickness of 250 µm. The slices were then recovered in 36°C NMDG-ACSF solution for 40 minutes, with gradual introduction of Na^+^ into solution following the Na^+^ spike-in procedures described by [26]. The slices were then transferred to room temperature HEPES-based (same as NMDG-prep solution, with 92 mM NaCl instead of NMDG, 2 mM MgSO_4_, 2 mM CaCl_2_) storage solution for the day.

#### 2.2.2 Patch-clamp recordings

Patch-clamp experiments were carried out with CleverExplore Neuroscience Workstation (MCI Neuroscience), including 10×air (CP-Achromatic, Zeiss) and 40×water-immersion (LUMPlanFL, Nikon) objectives, DIC optics, digital camera (MCI-CXE-B013-U), and EPC-10 USB Double amplifier (HEKA Electronics) using PatchMaster v. 2×92. The recording ACSF contained (in mM): 2.5 KCl, 1.25 NaH_2_PO_4_, 25 NaHCO_3_, 17 glucose, 125 NaCl, 2 CaCl_2_, and 1 MgCl_2_, feeding at a rate of 2.5 mL*/*min. The recordings were made at the average temperature of 29.5 ± 1.5 °C. The internal solution of glass pipettes (GB150F-10, Science Products, 5.5−9 MΩ) contained (in mM): 130 K-gluconate, 5 KCl, 10 HEPES, 0.6 EGTA, 2.5 MgCl_2_, 10 phosphocreatine, 4 ATP– Mg and 0.4 GTP– Na_3_.

Electrophysiological data was collected using `Patchmaster` software (HEKA Electronics) in whole-cell mode. After membrane breaking-in, cells were left to stabilize without current injection for 3 minutes. After this period, the spontaneous activity was recorded for at least 1 minute in the current-clamp configuration. Following that, additional current clamp recordings were carried out during which the neuron responses to injected current was evaluated. The current was injected in 11 steps, with the initial step bringing the cell to approximately −80 mV and the final step to approximately −40 mV. Cells with membrane potential under −40 mV were excluded from analysis. The collected data was converted to `abf` format and analyzed using `pClamp 10.7` (Molecular Devices) and `Matlab` (MathWorks).

### 2.3 Analysis

#### 2.3.1 Spike detection

Spike detection was carried out using the `Clampfit` software with threshold detection at 0 mV.

#### 2.3.2 Burst detection and classification

Bursts were detected using a simplified version of the *Poisson Surprise Method* [18] that defines bursts as unexpectedly short inter-spike intervals (ISIs) considering the average spiking frequency.

Assuming a Poisson distribution we calculated cumulative probability of observing one event for each ISI using the following formula:

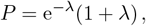

where *λ* is the product of the mean firing rate by the ISI. Hence, the Surprise index *S* was calculated as:

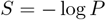

to mark the beginning of the burst, we set up a threshold of *P* ≤ 0.005 (*S* = 2.3). Since bursts are defined by an abrupt increase in spiking frequency that attenuates, we set up a lower threshold to define the end of a burst at twice the basal frequency (*S* = 1.045). We limited our analysis to bursts with 5 or more spikes (4 or more ISIs) in all cases.

Bursts evoked by hyperpolarizing steps were delimited and evaluated using the same approach. The only difference was that the baseline for the first post-hyperpolarization burst was estimated from a 500 ms segment before the beginning of the hyperpolarizing step. Only the first burst after each hyperpolarizing step was evaluated. Hyperpolarizing steps lasted one second and time between any two hyperpolarizing steps was 15.5 s.

Inspection of the data hints at the presence of square-wave, triangular and parabolic burst types. These burst types are differentiated by two factors: square-wave bursts have a raised baseline in comparison to quiescent interburst epochs (plateau), and feature a monotonic decrease in the spiking frequency (*i*.*e*., shortest ISI at the beginning and monotonically increasing along the burst), (Fig. 1 A, B) [17]). Parabolic bursts are characterized by the absence of a plateau and an increase in spiking frequency towards the middle of the burst (Fig. 1 E, F) [17]. Triangular bursts exhibit mixed features, lacking a plateau and displaying intraburst frequency patterns that span a continuum between monotonically decreasing and parabolic profiles (Fig. 1 C, D). Hence, to describe these bursts features, we selected three parameters. First, we estimated the presence of a burst plateau, measuring the difference between the burst minimum and the baseline from 300 to 30 ms before the first spike (Fig. 1 A, B). Any action potentials found within that window were manually excluded. In addition, to evaluate the monotonic decrease in the spiking frequency we computed two parameters. First, the relative position (in percentage) of the smallest ISI within the burst (Fig. 1) and second, the goodness of the fit (*R*^2^) of the burst ISIs to a single exponential function (Fig. 1 B, D, F):

**Figure 1:**
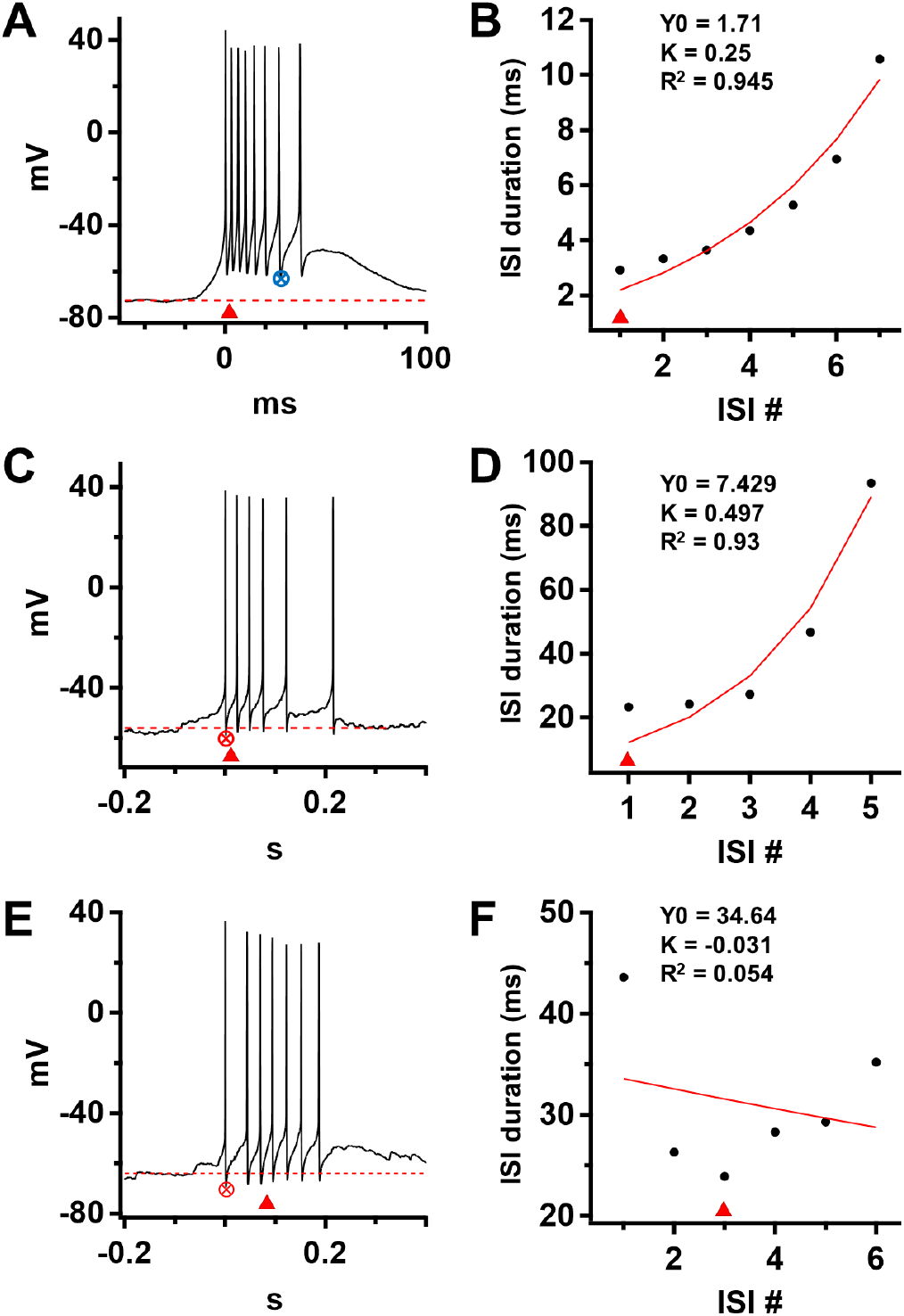
Parameters used for burst classification. **A, B**: case of a square-wave burst. The burst is on a plateau compared to the baseline, and has monotonically increasing ISIs. Hence it has the shortest ISI at the beginning of the burst and ISIs have a good fit to an exponential function (*R*^2^ = 0.945). **C, D**: case of a triangular burst, without a plateau during the burst and monotonically increasing ISIs. **E, F**: case of a parabolic burst, with no plateau during the burst and with ISIs arranged in approximately parabolic-shape, namely, with ISIs first decreaseing, then increasing, and minimal ISI being around the middle of the burst. This rather parabolic-like distribution of ISIs has a poor exponential fitting. Crossed circle indicates intra-burst minimum above (in blue) or below (in red) pre-burst baseline. The red dashed lines are the pre-burst baselines. Red triangle indicates shortest ISI in a burst. Red lines in **B, D, F** are exponential fitting on ISI index to ISI duration.

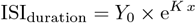

where *x* is the index of the ISI within the burst

#### 2.3.3 Statistics

Statistical analysis was carried out using `RStudio` v.2023.12.0 Build 369. The Shapiro-Wilk test was used to determine data normality, Levene’s test was used to determine homogeneity of variance. For comparisons, in case of normally distributed samples with equal variance a two samples unpaired t-test was used. In case one or more of variables being non-normally distributed a two samples Wilcoxon test was used.

### 2.4 Computational model

Based on the experimental data we developed a model reflecting the presence of two main types of bursting patterns together with a third one that is intermediate between the other two. The right panels of Fig. 2 show the outputs of the model corresponding to each of these three bursting patterns, namely square-wave (top), parabolic (center) and mixed-type or triangular (bottom); the left panels show recordings displaying the three different bursting behaviors. To identify these different bursting patterns in the computational model, we follow the standard approach introduced by J. Rinzel [24], termed *slow-fast dissection*, and apply it to an extended version of a classical threedimensional phenomenological bursting model called the Hindmarsh-Rose (HR) model [14]. The initial model from [14] has two fast variables, which can be associated with membrane potential and fast ionic dynamics, as well as one slow variable associated with slow ionic dynamics. We extend the model by adding a second slow variable, which can be related to an additional slow current, in order to capture parabolic bursting [10, 24].

**Figure 2:**
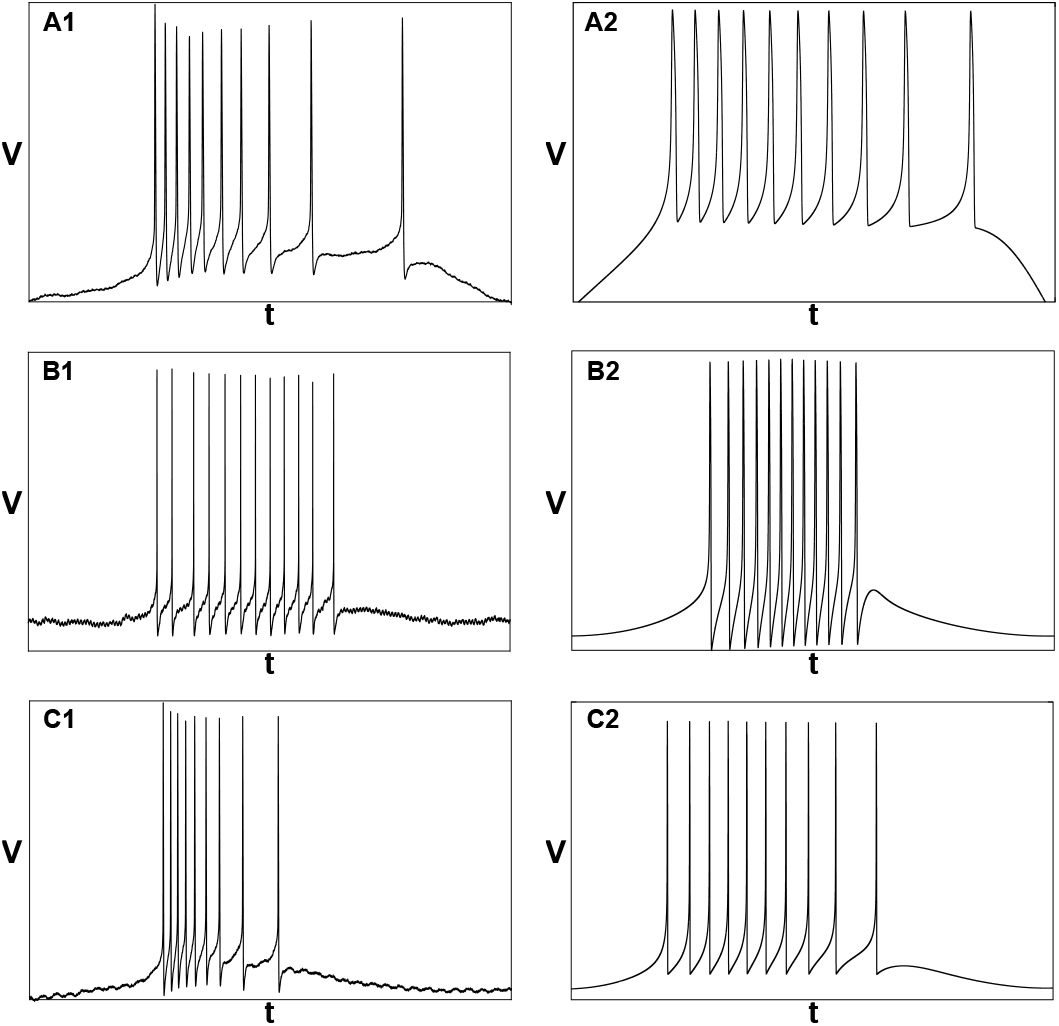
Experimental data (left panels) vs. mathematical model (1) (right panels), showing three bursting patterns identified in LHb neurons and captured by our idealised model. Namely: **A1**-**A2** square-wave bursting; **B1**-**B2** parabolic bursting; **C1**-**C2** triangular (mixed) bursting, at the transition between the other two patterns (see also Fig. 6). Fixed parameter values are: *a* = 0.08, *c* = 3, *d* = 1.8, *ε* = 0.01, *s* = 0.01, *x*1 = *−*3. Case-specific parameter values are: **A2** *b*0 = 0.52, *z*0 = *−*0.7163, *α* = *−*0.1; **B2** *b*0 = 1.5, *z*0 = *−*1.75, *α* = *−*0.1; **C2** *b*0 = 0.75, *z*0 = *−*2, *α* = 0.003.

In the case of models with two slow variables, the slow-fast dissection method consists in relating the bursting pattern produced by the complete model, projected onto a plane formed by the slow variables, with the two-parameter bifurcation diagram of the so-called *fast subsystem* obtained when freezing the dynamics of the slow variable and considering them as parameters in the fast differential equations. That particular bifurcation structure is the signature of the corresponding patterns, it allows to characterize it and, hence, to classify bursting patterns [11, 17, 24]. More precisely, one only needs two bifurcations of the fast subsystem in order to fully characterize a given bursting patterns, namely one bifurcation that organizes the transition between quiescence and burst in the complete model, and another bifurcation organizing the reverse transition.

According to the classification system introduced by Rinzel, square-wave bursting is associated with a saddle-node bifurcation of equilibria, initiating the burst, and a saddle homoclinic bifurcation terminating it. Note that this bursting pattern requires only one slow variable but it can also be obtained in a model with two slow variables. In contrast, parabolic bursting requires minimally two slow variables, and it corresponds to a bursting case in which both the initiation and termination of the burst in the full system are caused by the same bifurcation in the fast subsystem, namely, a saddle-node on invariant circle (SNIC) bifurcation. To be more precise, parabolic bursting is associated with a family of SNIC bifurcations in the fast subsystem, depending on two parameters. Hence, the full system has minimally two fast and two slow variables, unlike minimal square-wave bursting models, which have only one slow variable. Therefore, for the original threedimensional HR model (with one slow variable) to sustain parabolic-bursting activity, we need to augment it by one differential equation, which will correspond to an additional slow ionic variable. As mentioned above, such a four-dimensional model can also sustain square-wave bursting patterns, single saddle-node and homoclinic bifurcations of the fast subsystem being replaced by curves of such bifurcations.

Once we have a common computational model for both bursting patterns, the transition from one pattern to the other is organized in parameter space by a codimension-2 bifurcation point, namely a *saddle-node homoclinic (SNH)* point, also referred to as *saddle-node separatrix loop bifurcation* [17], in the vicinity of which one can generate bursting patterns of parabolic and square-wave types, respectively. At the boundary between these two bursting regimes, that is, in the vicinity of the SNH point in the plane of slow variables, one can obtain an intermediate bursting pattern, termed triangular or mixed-type, whose output looks like square-wave bursting for the initiation of the burst, and like parabolic bursting for the termination of the burst. The quiescent phase of mixed-type bursting trajectories has intermediate characteristics between the other two patterns; see Section 3 and Fig. 6 for details.

#### 2.4.1 Deterministic and stochastic versions of the model’s equation

The extended HR model used in the present work is composed by a set of four nonlinear differential equations, and it is deterministic. Its equations read:

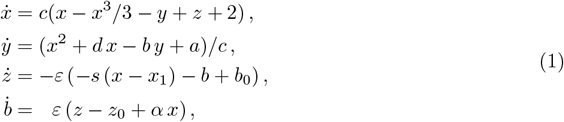

the last two equations corresponding to a slow dynamics on (*z, b*) as a phenomenological way to include voltage-gated slow channel dynamics. The parameter values are fixed in all three bursting cases, except for parameters *b*0, *z*0 and *α*, which allow to switch between cases; see the caption of Fig. 2.

We also consider a stochastic version of System (1) by adding noise terms to the two slow equations, as an elementary way to mimic channel noise. The resulting system of stochastic differential equations (SDEs) reads:

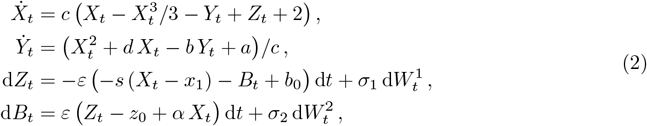

where 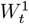 and 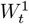 are two independent and standard Brownian motions, also independent from the initial conditions of the system.

## 3 Results

### 3.1 Experimental results

To characterize bursting activity of LHb neurons we performed patch-clamp recordings in currentclamp configuration of 61 LHb neurons, of which 41 showed spontaneous spiking activity. Following previous publications, we classified their activity according to the spiking patterns into regular, non-regular, and bursting [27]. To identify and define bursts, we followed a statistical criterion based on the *Poisson Surprise method* (see section 2.3.2 of Methods). Using this criterion and in agreement with previous reports [3, 9, 22] only 15 cells spontaneously displayed bursting activity (Fig. 3 A).

**Figure 3:**
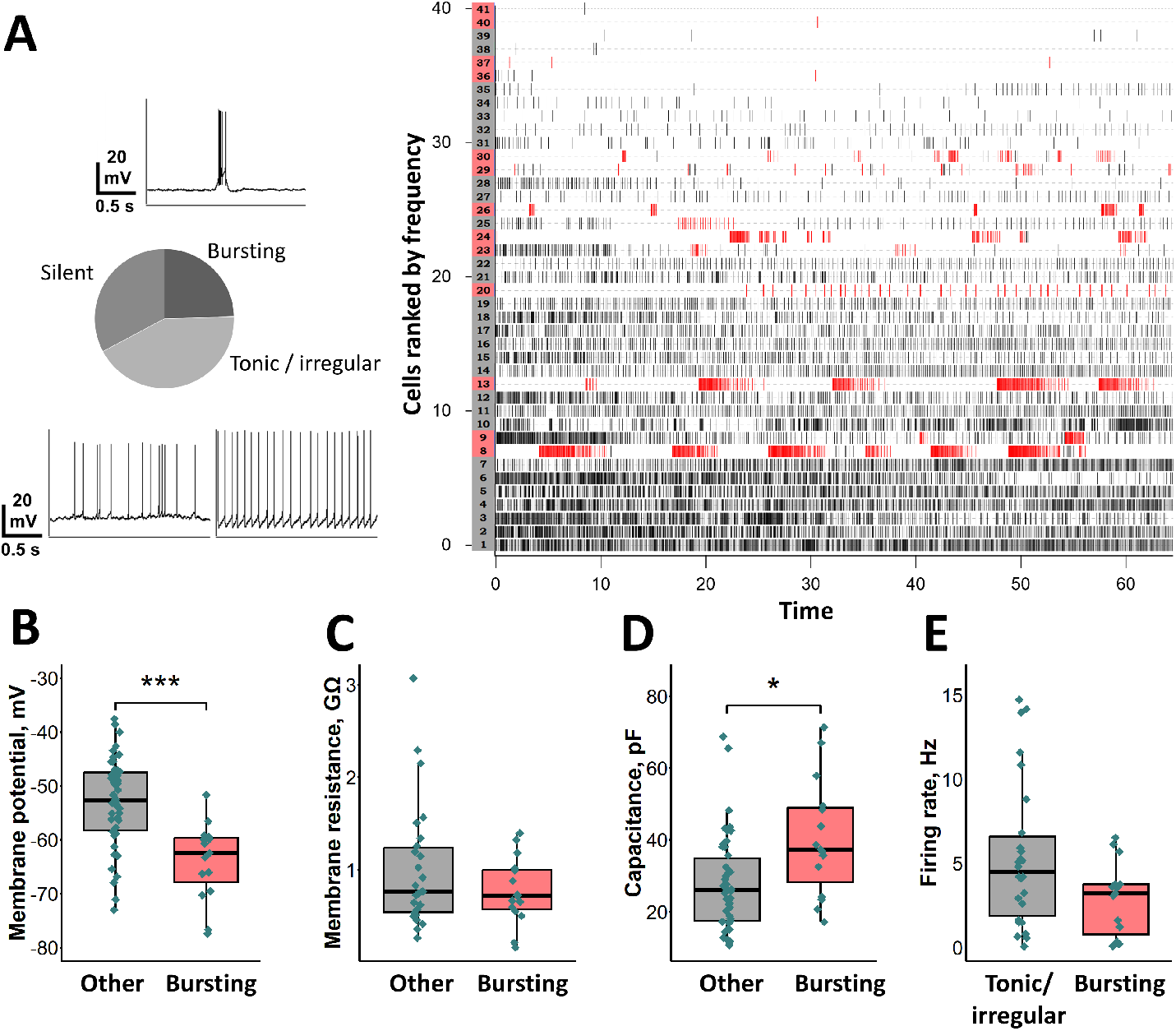
LHb neurons parameters and classification of spontaneous activity. **A**, cell classification by spontaneous activity; 15 bursting cells, 26 tonic/irregular firing, and 20 silent. Above and below the pie chart: examples of cell activity. Right: raster plot of action potentials in cells ranked by mean spiking frequency. Bursting cells and individual bursts are highlighted in red. Two cells that displayed only one burst are not highlighted. **B**, membrane potential; **C**, membrane resistance and **D**, membrane capacitance in bursting and other cells (including silent, *n* = 61). **E**, firing rate during the first 60 s of the recording, excluding silent cells (non-bursting *n* = 41).

Bursting cells were characterized by a more negative resting potential (−65.25 ± 0.66 mV for bursting neurons compared with −53.13 ± 8.35 mV for non-bursting neurons; *p* = 0.000 042 3 Student’s t-test; *N* = 15 and 46 respectively, Fig. 3 C), and higher capacitance (40.25 ± 16.32 pF vs. 27.72 ± 13.06 pF; *p* = 0.00626, Wilcox test, Fig. 3 D) and did not show differences in membrane resistance (0.77 ± 0.36 GΩ vs. 0.87 0. ± 67 GΩ for bursting and non-bursting neurons, *p* = 0.722, Wilcox test, Fig. 3 E). In addition, the mean spiking frequency (excluding silent cells) was not significantly different between bursting and non-bursting neurons (2.87 ± 2.19 Hz vs. 5.42 ± 4.39 Hz for bursting and non-bursting cells, *p* = 0.0673 Wilcox test, Fig. 3 F), supporting the notion that at the LHb bursting activity does not represent an increase in the average output.

Visual inspection of spontaneous bursts revealed the presence of different types of bursts, including square-wave, parabolic, and triangular. Square-wave and parabolic bursts are distinguished by their membrane potential dynamics and the evolution of spiking frequency within the burst. Square-wave bursts are characterized by a sustained depolarized plateau and a monotonic decrease in spiking frequency (Fig. 1 A), whereas parabolic bursts lack a plateau and exhibit a peak in spiking frequency near the middle of the burst (Fig. 1 C). Triangular bursts represent a transitional form, lacking a clear plateau and displaying an intraburst frequency profile that bridges the properties of square-wave and parabolic bursts.

To quantify these features, we calculated three parameters from the 809 bursts recorded in 15 spontaneously bursting neurons: the minimum voltage within the burst, the goodness of fit of the ISIs to an exponential decay (*R*^2^), and the relative position of the minimal ISI within the burst (see Methods). Fig. 4 displays these results for all bursts and individualized by cell.

**Figure 4:**
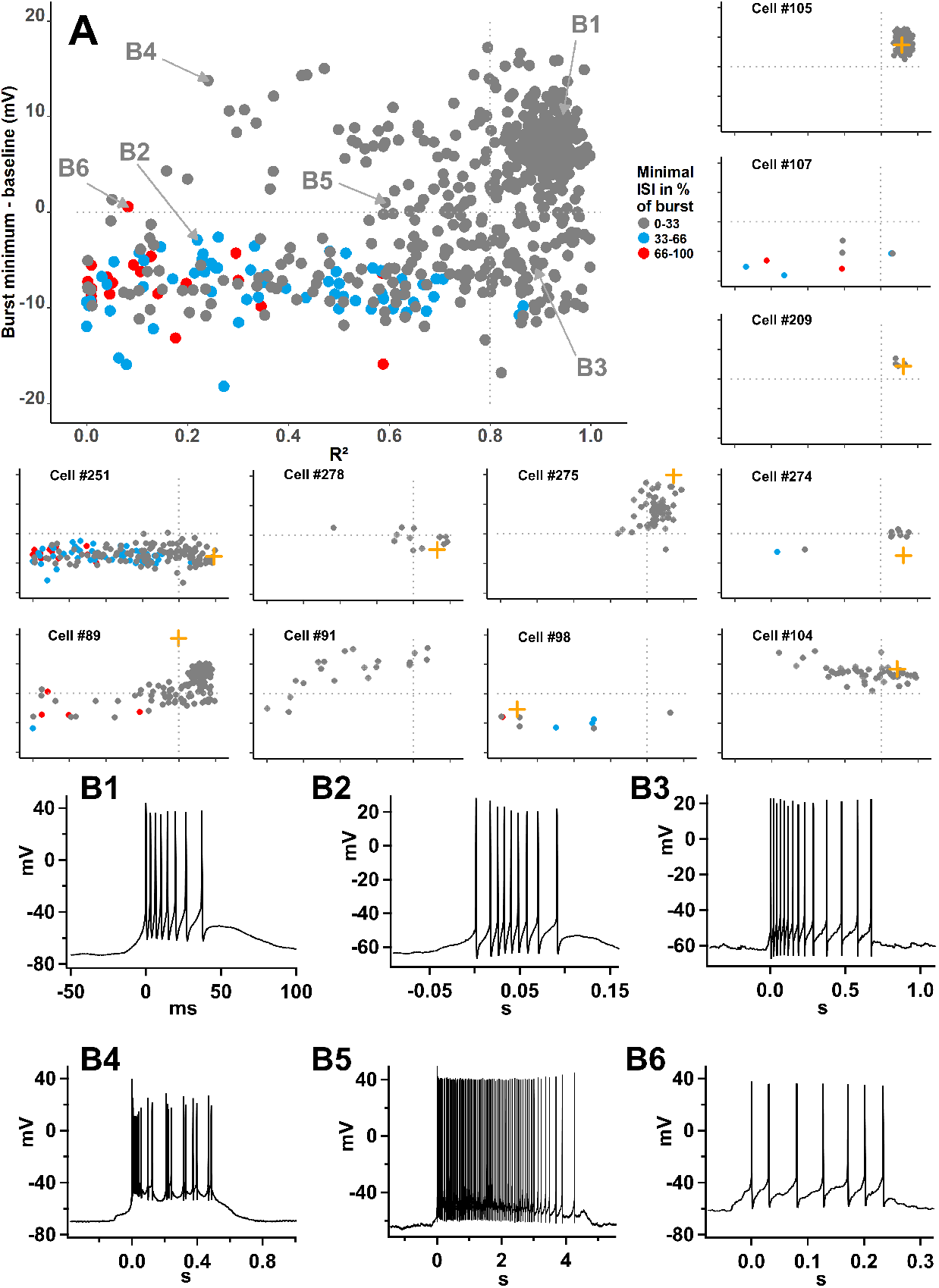
Parameters of spontaneous bursts. **A:** The central graph shows the distribution of 809 spontaneous bursts recorded from 15 cells, while the smaller lateral graphs present representative examples from individual cells. A distinct cluster of bursts appears in the upper-right region of the central plot, characterized by square-wave-like features: a plateau phase (Burst minimum *<* 0), good exponential fit of the ISIs (*R*^2^ *>* 0.8), and a minimal ISI at the onset of the burst (indicated by black dots). Panel **B1** shows a representative example of a burst from this region. Notably, this burst pattern is dominant in cell 275 and uniquely observed in cells 105 and 209. In the lower half of the graph (Burst minimum *<* 0), bursts are distributed across a spectrum. On the left side, parabolic-type bursts show poor exponential fit and a minimum ISI occurring mid-burst (**B2**). Towards the right side, triangular transitional bursts are distributed on a gradient of increasing *R*^2^ values along with an increasing proportion of bursts with the minimum ISI at the beginning and minimum ISIs (exemplified in **B3**). Panels **B4**–**B6** show examples of other burst types not fitting these main categories: irregular bursts with a plateau but without monotonic decay (**B4**); long bursts that begin and end with monotonic ISI decay but include a second long high-frequency spiking plateau phase sustained by depolarization (**B5**); and sparsely observed bursts with an inverted parabolic shape, where the lowest spiking frequency, corresponding to the longest ISI, occurs in the middle of the burst (**B6**). Dashed horizontal line: burst minimun = 0. Dashed vertical line: *R*^2^ = 0.8. For the sake of clarity only cells with more than 5 bursts are individually presented. Orange crosses represent the burst evoked in the same cells by a 1 s hyperpolarizing step from ≈ −80 mV (see Fig. 5 for details). Cell 107 became tonic after the hyperpolarizing step, whereas cell 91 generated a burst consisting of only two spikes.

**Figure 5:**
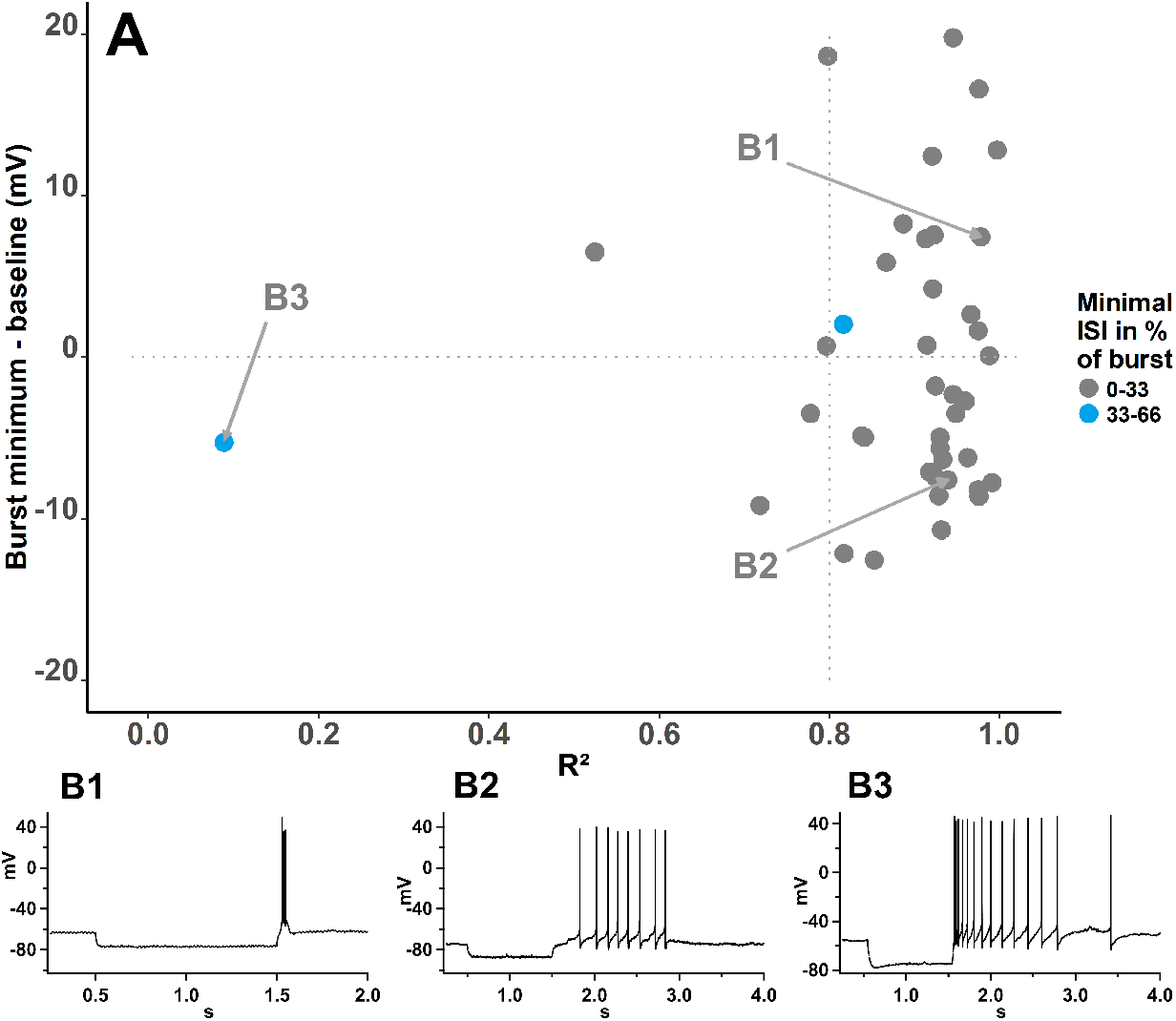
Parameters of hyperpolarization evoked bursts recorded from 41 cells in LHb. Bursts were evoked by a hyperpolarizing current step that brings the membrane potential to ≈ −80 mV. Only bursts of at least 5 spikes were included. Each dot is the first burst after the end of the step. Burst type prevalence by quadrants and other constraints are same as in Fig. 4. Baseline for plateau evaluation is taken from the period before the hyperpolarizing step.

A large proportion of the recorded bursts, located in the upper right quadrant of Fig. 4 A main panel, exhibit square-wave burst features: a plateau (burst minimum *>* 0), a good exponential fit of the ISIs (*R*^2^ *>* 0.8), and the shortest ISI located at the beginning of the burst (grey dots, Fig. 4 B1). Moreover, 3 out of 15 cells (cells: 105, 209 and 275) almost exclusively exhibited this type of bursting activity, (Fig. 4 A).

In the lower half of Fig. 4 A (burst minimum *<* 0), bursts are distributed on a gradient. At the left extreme, we found bursts that lacked a plateau, showed poor exponential fitting of the ISIs, and had their minimal ISIs near the middle of the burst (blue dots, Fig. 4 B2), features consistent with parabolic bursts (see Fig. 1). Also, few bursts in this quadrant exhibited their minimal ISIs toward the end of the burst, which does not conform to the parabolic burst profile (red dots, Fig. 4, example in B6). In the lower half of Fig. 4 A, towards the right side, along with the goodness of the ISI exponential fitting, also the proportion of bursts with the minimum ISI a the beginning of the burst increases and, at the right extreme, there is a dense population of bursts with a good ISI exponential fitting (*R*^2^ *>* 0.8) and minimum ISI at the beginning of the burst, indicative of a monotonic decay. This population of events, characterized by features transitioning between parabolic and square-wave type bursts, represents the triangular burst type (exemplified in Fig. 4 B3).

Finally, bursts located in the left part of the upper half of Fig. 4 exhibited a plateau and had their minimal ISIs at the beginning of the burst, but did not fit well to a single exponential decay. These bursts may represent more complex dynamics that do not clearly fit into any of the three defined categories (see Fig. 4 B4 for an example). Additionally, within the upper half of the graph (burst minimum ≥ 0), we observed long bursts characterized by square-wave-like decreases in spiking frequency at both the onset and offset, separated by a prolonged period of nearly constant firing rate that could last several seconds (Fig. 4 B5). This unusual structure led to variable *R*^2^ values, making these long bursts not visually identifiable in Fig. 4.

The majority of the spontaneously bursting cells displayed more than one type of burst, indicating that different bursting patterns could be generated by the same LHb neuron and represent different dynamic states rather than different cell types.

It has been consistently reported that the majority of LHb neurons generate burst of action pontential upon hyperpolarization. We therefore extended our analysis to hyperpolarization-triggered bursts. Most LHb neurons (41/61) exhibited rebound bursting activity. To compare hyperpolarization triggered and spontaneously bursting we measured the same parameters of spontaneous bursts on the first burst after a hyperpolarization step of ≈ −80 mV. Hyperpolarization-induced bursts presented predominantly features of square-wave or triangular, with only a few bursts showing parabolic features. This observation suggests that hyperpolarization-triggered bursting represents a fraction of the variety of bursting behaviors LHb neurons could display. Indeed, the hyperpolarization protocol carried out on cells that also had spontaneous bursting tended to produce bursts with higher *R*^2^ values than that of most spontaneous bursts of the same cell (Fig. 4, orange crosses in individual cell panels).

From the neurodynamics perspective square-wave and parabolic bursts represent different phenomena. Since individual LHb neurons could display those two patterns of activity and/or the intermediate triangular bursts, in the next section, we develop a model that could account for that variety of behaviors.

### 3.2 Modelling results

System (1) has explicit timescale separation with two fast variables *x* and *y*, which mimic membrane potential and fast channel dynamics, respectively, and two slow variables *z* and *b*, which mimic slow channel dynamics. Standard slow-fast analysis consists in setting *ε* = 0 in System (1), which yields the so-called fast subsystem. In the fast subsystem, the slow variables become parameters and the main interest in studying the fast subsystem comes through its bifurcation structure with respect to these distinguished parameters. Indeed, when reinstating *ε* = 0, one realizes through numerical simulations (and can prove mathematically) that the full system dynamics remain most of the time close to families of attractors (stationary or periodic) of the fast subsystem. When the system contains two slow variables, and hence the fast subsystem two distinguished parameters, the corresponding bifurcation diagram of interest is naturally a two-parameter bifurcation diagram, onto which it is useful and customary to superimpose solutions from the full system, projected onto this diagram; see Fig. 6. This approach is particularly suited to studying bursting dynamics in the full system, in which case the projection onto the plane of slow variables typically shows a slow path flowing across bifurcation curves of the fast subsystem. In particular, bifurcation of equilibria, *e*.*g*., saddle-node (SN) bifurcations and Hopf (HB) bifurcations, and bifurcation of limit cycles, *e*.*g*., saddle homoclinic (Hom) bifurcations and SNIC bifurcations.

**Figure 6:**
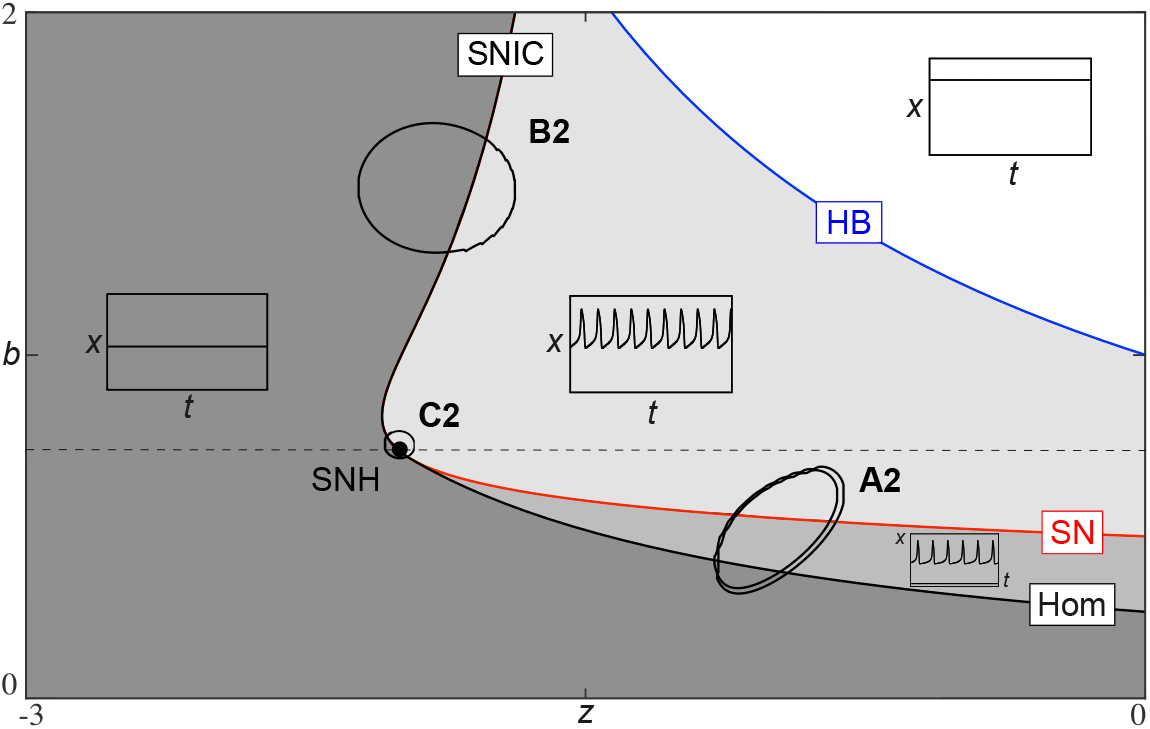
Two-parameter bifurcation diagram of the model’s fast subsystems (*i*.*e*., when the slow variables’ dynamics are frozen and replaced by parameters) explains the two main bursting patterns found, as well as, the intermediate one. The red curve correspond to a family of saddle-node bifurcations of equilibria labelled SN, the black curve corresponds to a family of homoclinic bifurcations, and the blue curve to a family of Hopf bifurcations. The first two curves meet at a codimension-2 bifurcation point called *saddle-node homoclinic* and labelled SNH. Above that point, in the two-parameter plane, the two curve coincide and correspond to a family of saddle-node on an invariant circle (labelled SNIC) bifurcation. Below SNH, the black curve corresponds to a family of saddle-homoclinic bifurcations labelled Hom. Round-shaped curves correspond to the slow evolution of parameters *z* and *b*. When the values of *b* stay below the SNH point, then the bursting pattern observed in the full system is of square-wave type and corresponds to the profile A2 from Fig. 2. In contrast, if the value of *b* is located above the SNH point, then the observed bursting pattern in the full system is of parabolic type and it corresponds to the profile B2 of Fig. 2. At the transition from one pattern to the other, we have trajectories of the full system that revolve around the SNH point, giving triangular (or mixed) bursting orbits like the one shown here, which corresponds to the profile C2 of Fig. 2.

In Fig. 6, on top of the two-parameter bifurcation diagram System (1)’s fast subsystem with respect to *z* and *b*, are superimposed the three bursting trajectories shown in Fig. 2 A2, B2 and C2, respectively, projected onto the (*z, b*) plane. As explained above, this plotting strategy allows to shed important light onto the different bursting patterns depending on the region of parameter space visited by the slow dynamics. In the present case, the pattern corresponding to panel A2 in Fig. 2 is of parabolic bursting type, the pattern corresponding to panel B2 is of square-wave type, and the intermediate pattern corresponding to panel C2 is of mixed type and indeed it revolves around the SNH point. All three patterns can be associated with the crossings of different bifurcation curves of the fast subsystem. Namely, parabolic bursting occurs when the slow variables drive the dynamics back and forth across a curve of SNIC bifurcations (black curve above the SNH point on Fig. 6), while square-wave bursting is the result of a slow passage through a curve of saddle-node bifurcations of equilibria (red curve) and a curve of saddle-homoclinic bifurcations (black curve below the SNH point). The mixed-type (triangular) bursting pattern corresponds to an initiation of the burst as the trajectory crosses the SN curve, and a termination of the burst as it crosses the SNIC curve, hence giving a mix of both main patterns. Square-wave and parabolic bursting patterns dominate and, loosely speaking, they correspond to cases where the slow variables evolve below and above, respectively, the SNH point; see the dashed line in Fig. 6. In contrast, the mixed-type bursting pattern is obtained from trajectories whose slow dynamics revolve around the SNH point.

The effect of noise on this two-parameter structure allows a switch from one pattern to the other one following a slow random path. This can produce time series where both bursting patterns are successively present Fig. 7 reproducing the behavior observed in LHb neurons.

**Figure 7:**
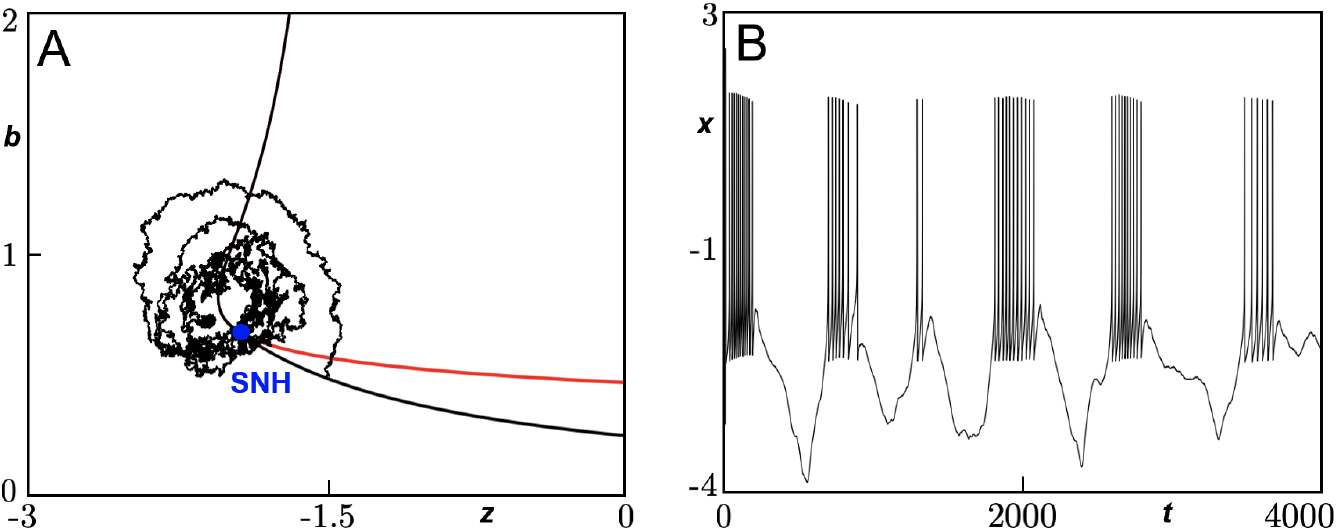
Simulation of the extended HR model with added stochastic input on the slow dynamics, System (2), leading to mixed bursting patterns. Fixed parameter values are as in Fig. 2, the other parameter values being: *b*0 = 0.8, *z*0 = *−*1.8, *α* = *−*0.1, *σz* = *σb* = 0.75.

## 4 Discussion

In the last decades, the LHb has been increasingly recognized as a central hub for the neural processing of aversive stimuli [15, 21, 23]. This conceptualization is supported by three observations consistently reported across models. First, the LHb exhibits systematic excitation in response to aversive stimuli. Second, its activity increases in states characterized by heightened aversiveness, such as stress or drug withdrawal. Third, its activation is sufficient to trigger aversion [15].

Physiologically, the LHb is characterized by low intraconnectivity and a predominance of glutamatergic neurons. Thus, its output is primarily shaped by the intrinsic properties of individual neurons and their synaptic interactions and not by intra-LHb interactions.

Both in vivo and ex vivo, LHb neurons exhibit high basal activity and a diverse spiking repertoire, which can be classified as silent, tonic, irregular, or bursting [28]. Surprisingly, no consistent relationship has been found between these spiking patterns and anatomical localization, morphology, or gene expression [13, 28]. In fact, most LHb neurons can express all four spiking modes depending on membrane potential [9, 28]. Bursting activity, in particular is promoted by membrane hyperpolarization [9, 28]. Accordingly, we found spontaneously bursting LHb neurons to be more hyperpolarized than silent or tonically active neurons.

Ex vivo, only a minority of LHb neurons exhibit spontaneous bursting activity. However, animal exposure to chronic stressors persistently increases the proportion of bursting neurons. This enhancement in LHb bursting has been specifically linked to the emergence of depressive-like phenotypes in mice [3, 6, 7, 8, 9, 16, 22, 25, 30]. Remarkably, ketamine has been shown to reverse these behavioral alterations by suppressing bursting activity in the LHb [5, 20, 30].

Despite its importance, bursting activity in the LHb has been characterized only in broad terms, often described as periods of elevated activity defined by absolute frequency criteria, or in some cases left undefined [6, 8, 9, 16, 22, 25, 30], but see [3, 7]. This study represents a step toward addressing this gap by aligning LHb bursting activity with the systematic classification frameworks developed for bursting in other neuronal contexts [11, 17].

We observed that LHb neurons spontaneously exhibited a wide variety of bursting patterns. These ranged from square-wave-like bursts, characterized by a plateau and monotonic decrease in spiking frequency, to parabolic-like bursts, which lack a plateau and display peak frequency in the middle of the burst. Between these distinct types, we also identified a substantial proportion of transitional triangular-like bursts, that is, bursts without a plateau in which the intraburst frequency varies from pure monotonic to close to parabolic. Additionally, we detected other bursts that did not fit into these categories, among them, long bursts lasting several seconds, displaying square-wave-like features at the beginning of the burst, triangular-like end, and maintaining a relatively constant high spiking frequency in the middle. Notably, these bursts have been recently linked to neuron-to-glia communication [29, 4].

To expand the number of cells in which we could study bursting activity we applied hyperpolarizing steps, a manipulation widely shown to induce rebound bursting. However, these rebound bursts were almost exclusively of the square-wave-like type, suggesting that hyperpolarization promotes only a subset of the potential bursting modes of LHb neurons.

The ionic conductances underlying square-wave-like bursts induced by hyperpolarization have been well described by Yang et al. [30] and depend on low-threshold voltage-dependent calcium channels and, to a lesser extent, HCN channels. In addition, long bursts have also been shown to depend on cyclic nucleotide-gated channels and neuron–oligodendrocyte coupling [4, 29]. On the other hand, the ionic mechanisms underlying parabolic and triangular bursts in the LHb remain unknown and are less well characterized in other systems, where they may involve persistent sodium channels and M-type potassium channels [2, 12].

In our experiments, LHb bursting did not correspond to an increase in mean neuronal spiking frequency, a finding recently supported in vivo by other authors [31]. This suggests that the link between LHb bursting and negative motivational value may arise from the dynamics of bursting itself rather than by the average LHb output. In this context, square-wave and parabolic bursts, could represent different output modes with different impact at the presynaptic terminals governed by their own facilitating or depressing dynamics. The former, characterized by an initial highfrequency spike followed by a decay, would be amplified by facilitating synapses and filtered by the depressing ones. The latter, with low initial frequencies that peak toward the middle of the spike train, would exhibit the opposite profile.

Interestingly, most spontaneously bursting neurons exhibit more than one burst type, indicating that these patterns are not limited to specific cell classes. To account for these complex dynamics, we developed an idealized model (System (1)) that is capable of displaying each of these patterns separately based on slightly different values of two key parameters, the boundary between them being displayed in parameter space by the presence of a codimension-2 bifurcation point, which we identify as SNH. We envisage these two parameters to be related to slow ionic conductances in real neurons. Hence, simulating the model with added stochastic components onto the differential equations where these two parameters appear (System (2)) allows to produce simulations displaying more than one burst type.

The model predicts that LHb neurons can transition between distinct bursting modes-ranging from square-wave to parabolic bursting, when shifted around the SNH critical point in a space likely governed by slow ionic conductances. Hence, in biological neurons, it is reasonable to speculate that sustained changes in these putative slow conductances could bias the system toward one bursting mode over the other, as seen in the case of the shift in membrane potential mediated by astrocytic Kir 1.4, which favors square-wave bursting [9]. Synaptic inputs may introduce the stochastic fluctuations necessary to drive transitions across this equilibrium point what would also explain the great amount of evidence linking synaptic changes at the LHb with depressive-like behaviors [23].

Hence, our formalism predicts that the bursting mode predominant on LHb neurons is related slow conductances balance, and not to a single conductances. This, highlights the multifactorial nature of neuronal dynamic behavior, and is particular relevant for translational approaches aimed at controlling LHb bursting activity. Future experimental and modeling studies will be essential to determine how these different bursting modes influence mood and aversion, and how the interplay between intrinsic and synaptic conductances regulates their balance.

## Data Availability

All original data from which graphical data is generated is archived and fully available to The Journal upon request.

## Acknowledgments

DF, ESG, SR and JP acknowledge support of SILICON BURMUIN no. KK-2023/00090 funded by the Basque Government through ELKARTEK Programme. ESG acknowledges support from the grant PID2021-125763NB-I00 funded by the MICIU / AEI and from the Fundacion Tatiana, SR is supported by the grant PID2023-146683OB-100 funded by MICIU / AEI / 10.13039 / 501100011033 and by ERDF, EU. Additionally, he is supported by Ikerbasque Foundation and the Basque Government through the BERC 2022-2025 program and by the Ministry of Science and Innovation: BCAM Severo Ochoa accreditation CEX2021-001142-S / MICIU / AEI / 10.13039/501100011033. Moreover, SR acknowledges the financial support received from BCAM-IKUR, funded by the Basque Government by the IKUR Strategy and by the European Union NextGenerationEU/PRTR. JP acknowledges support by the Ikerbasque Basque Foundation for Science, EU COFUND H2020-MSCA-COFUND-2020-101034228-WOLFRAM2, and the Achucarro Basque Center for Neuroscience.

